# Identification of a conserved RNA structure in the *TNFRSF1A* 3’UTR: Implications for post-transcriptional regulation

**DOI:** 10.1101/2025.06.18.660452

**Authors:** Van S. Tompkins, Warren B. Rouse, Jibo Wang, Michael E. Woodman, Ernst R. Dow, Theodore C. Jessop, Walter N. Moss

## Abstract

Tumor necrosis factor receptor superfamily 1A gene (*TNFRSF1A*) encodes the TNFR1 protein, a critical regulator of inflammation implicated in various diseases. Using `ScanFold` with the Integrative Genomics Viewer (IGV) GUI, we identified novel RNA structural elements within the *TNFRSF1A* gene. Focusing on the 3’UTR, these structures were characterized using reporter assays and targeted DMS-MaPseq. We identified a structured region that may play a role in regulating TNFR1 translation and that was also found to associate with HuR, a key regulatory RNA-binding protein. These findings provide a framework for identifying and characterizing potential functional RNA structures in therapeutically relevant genes, suggesting a new layer of post-transcriptional regulation for TNFR1 expression.

## 1. Introduction

The tumor necrosis factor receptor superfamily 1A gene (*TNFRSF1A*) encodes TNFR1 (tumor necrosis factor receptor 1), a transmembrane glycoprotein containing four cysteine rich extracellular domains that can be proteolytically cleaved to produce a soluble form of the receptor [1]. Both forms interact with its ligand, tumor necrosis factor alpha (TNF-*α*), a proinflammatory, pleiotropic cytokine produced by immune cells in response to a variety of stimuli. Binding of TNF-*α* to the extracellular domain of membrane bound TNFR1 activates signal transduction of different pathways through formation of TNFR1 homotrimers, stimulating interaction with other signaling molecules [2]. After signaling is complete, the complex is internalized and dissociates in the endosome [3]. If the receptor is proteolytically cleaved into its soluble form, it is released into the cytoplasm where it can act as an antagonist through interaction with free TNF-α [1]. When active, the receptor plays roles in cell survival and inflammation, apoptosis, or necroptosis. Mutations of this gene have been associated with TNF receptor-associated periodic syndrome (TRAPS) and multiple sclerosis in humans [4], whereas dysregulation of TNFR1 expression is observed in many associated disease states including rheumatoid arthritis and cancer [5–7]. A better understanding of the factors that control TNFR1 expression will provide functional and mechanistic insights with potential to provide routes for therapeutic modulation: e.g., via emerging small-molecule based approaches for targeting RNA structure [8–10]. RNA structure is one such factor that deserves more careful consideration.

RNA structure plays an important role in post-transcriptional regulation of mRNAs by adjusting access to primary binding regions, directing 3D spatial arrangement of cis-regulatory structures and trans-acting factors, and more. Unlike proteins, RNA folding is hierarchical with secondary structures forming before tertiary structures. Most of a RNA’s folding energy comes from hydrogen bonding of base pairing and base stacking interactions. Because the formation of RNA secondary structure largely determines what 3D structures are possible, the secondary structure is highly informative. Secondary structures are important to every step of RNA biology such as splicing, polyadenylation, localization, translation, and stability [11]. Similarly, structures across a pre-mRNA or transcript that are crucial for functions in cells can bind or facilitate the binding of proteins to RNA that may then execute functions [12, 13]. One such RNA binding protein is called HuR (encoded by *ELAVL1*). HuR binding predominantly stabilizes mRNA and can lead to increased translation [14, 15]. Finding these secondary structures and the proteins that bind them will help uncover the utility of mRNA as a therapeutic target.

To find RNA secondary structures with functional propensity, we previously developed `ScanFold` [16], a bioinformatics tool that employs a sliding window approach to predict local RNA secondary structure characteristics. `ScanFold` calculates important RNA folding metrics, including a thermodynamic z-score, which measures sequence-ordered thermodynamic stability. This z-score quantifies the difference in stability between the predicted minimum free energy (MFE) of a natively ordered RNA sequence and the MFE of randomized sequences with similar nucleotide content. Negative z-scores indicate that the native RNA sequence is more likely to fold into a specific, stable structure, suggesting potential functional significance. `ScanFold` leverages this thermodynamic bias to generate a weighted consensus structure z-score, highlighting recurring base pairs that contribute to highly stable and potentially functional RNA elements [17]. We also developed an implementation of the Integrative Genomics Viewer (IGV) that can visualize (and generate) `ScanFold` data to aid in RNA regulatory motif discovery through comparisons to other forms of complementary “omics” data [18]. These approaches were successfully used to discover functional, and targetable structures (via small molecules) in human genes, such as MYC [19, 20] and AR [21] as well as in SARS-CoV-2 [22]. Notably for AR [21] and for whole genome EBV-positive BJAB-B1 cells [20], `ScanFold` structure predictions have been enhanced using dimethyl sulfate (DMS) structure probing. This method enables in-cell snapshots of RNA structure through chemical accessibility [23].

In this study, we examine the RNA secondary structure of the *TNFRSF1A* gene, which encodes the TNF receptor 1 (TNFR1) protein. We first employed `ScanFold` to predict the RNA folding landscape of the *TNFRSF1A* gene and its MANE transcript (Matched Annotation between NCBI and ENSEMBL). We then focused our experimental analyses on the 3’ untranslated region (UTR), examining several regions in reporter assays. Further, we used both in vitro and in cell dimethyl sulfate (DMS) probing to map the RNA structure. This combined computational and experimental approach revealed functional RNA structures within the 3’UTR, including a structured element indicated both to influence *TNFRSF1A* expression and to associate with HuR.

## 2. Materials and methods

### 2.1. ScanFold

`ScanFold` is an RNA secondary structure prediction algorithm that identifies regions of local structural stability, unusual sequence-ordered stability, and likely functionality from long sequences. It achieves this by generating consensus structures whose base pairs are weighted by their contribution to ordered structural stability. In brief, `ScanFold` uses a scanning window to analyze RNA sequences of interest (here the *TNFRSF1A* gene and MANE transcript). Each window is folded by `RNAfold` [24] to calculate its native minimum free energy (MFE) and associated base paired secondary structure. The sequence of each window is shuffled, using mononucleotide shuffling, and folded 100 times to calculate an average randomized MFE value. These values are used to calculate the thermodynamic *z*-score. These z-scores are then used to generate a weighted consensus secondary structure model based on paired nucleotides that recur across low z-score analysis windows [16, 17]. A negative z-score indicates that the native sequence is ordered to form a more stable structure than other possible versions of the RNA with the same length and nucleotide composition. The consensus structures are biased towards sequence-ordered stability suggesting potential functionality via evolutionary preservation. All low z-score structures (≤ - 2) are extracted for further analysis. Metrics obtained from `ScanFold` include MFE, ΔG z-score, and ensemble diversity (ED) [16, 17]. In this study we used the stand-alone version of `ScanFold` to analyze the gene and transcript, and the integrated version, `IGV`-`ScanFold` [18], in a secondary analysis of the 3’UTR for ease of visualizing complementary datasets in the genomic context. For all `ScanFold` runs, we used a 120 nt window, a 1 nt step, 100 randomizations per window, mononucleotide shuffling, 37°C temperature, competition of 1 (to demand that only one unique base pair per nucleotide is possible), and extraction of structures with z-scores ≤ -2. All `ScanFold` output files can be found in **S1 File** (z-score, MFE, ED, predicted base pairs, etc.).

### 2.2. Cell culture

Three different human cell lines were used in this study: HeLa, Lenti-X, and THP1. HeLa (ISU Hybridoma Facility) and Lenti-X (Clontech) cells were maintained in DMEM, while THP1 cells (Eli Lilly) were maintained in RPMI-1640 at 37°C in 5% CO_2_. All media was supplemented with 10% FBS, 100 U/µg per mL penicillin/streptomycin, and 2 mM L-glutamine. All media, supplements, and Dulbecco’s phosphate buffered saline (DPBS) were obtained from Gibco/Thermo-Fisher. All cell lines were passaged at 80-90% confluence, used between passages 5-20, and regularly tested for mycoplasma [25].

### 2.3. In vitro RNA preparation for dimethyl sulfate (DMS) probing

*TNFRSF1A* sequences of interest (full-length 3’UTR, structure 7, and d7-deletion of structure 7 from 3’UTR) were cloned into a novel in vitro probing cassette (IVPC) within the pCR2.1 plasmid. The cassette contains three spacer hairpins (MFE between -11 and -12 kcal/mol) at the 5’ most end followed by an “insulator” hairpin (MFE of 14 kcal/mol). Adjacent to these hairpins, the 5’ and 3’ linker hairpins used by the Weeks Lab [26] were inserted to flank the MCS containing the XhoI restriction site for HiFi cloning. At the 3’ end of the cassette, adjacent to the 3’linker hairpin, three additional spacer hairpins (MFE between -11 and -12 kcal/mol) were also added (**S1 Fig**). This cassette was designed to aid in structure folding through insulation and to increase the sequence length for library preparation after probing. RNA was prepared from a total of 2 µg of each plasmid containing the sequences of interest. Plasmids were digested with SpeI (NEB) for run-off T7 in vitro transcription. Digests were validated via agarose gels, and the linearized plasmids were purified using a PCR cleanup kit (Zymo) following the manufacturer’s protocol. T7 in vitro transcription was done following a modified protocol. Here, 500 ng of purified linear DNA was incubated with 5x T7 buffer (Invitrogen), 10 mM NTPs (Invitrogen), 50 mM DTT (Invitrogen), and water for 1 hr at 37°C. After 1 hr T7 polymerase (Invitrogen) was added to the reaction and incubated for 1 hr at 37°C. After completion of the reaction, TURBO DNase (Invitrogen) was added to the reaction and incubated for 15 min at 37°C. The resulting in vitro transcribed RNA was purified using TRIzol® (Invitrogen) and Direct-zol Plus RNA mini-prep kit (Zymo) following manufacturer’s protocol. Purified RNA was heated to 75°C for 15 minutes and cooled to room temperature before 100 – 500 ng was used to verify the correct RNA product size via urea-PAGE (National Diagnostics Urea-Gel SequaGel) and SYBR Green II RNA Gel Stain (Thermo-Fisher) on a BioRad Gel Doc EZ Imager.

### 2.4. In vitro dimethyl sulfate (DMS) probing

In vitro DMS probing was done using a modified protocol from the Rouskin lab [27–29]. Briefly, 88 µl of refolding buffer (10 mM Tris pH 8.0, 100 mM NaCl, 6 mM MgCl_2_) was added to a 1.5 mL tube for each reaction and prewarmed to 37°C. A total of 10 picomoles of each RNA (1-4 µg) were added to a PCR tube and the final volume was brought to 10 µl with nuclease free water. Samples were denatured at 95°C for 1 minute and snap cooled by immediately placing on ice. All cooled RNA (10 µl) was added to the correct tube containing the refolding buffer at 37°C. Samples were gently pipette mixed and incubated for 15 minutes at 37°C. After the 15-minute incubation, 2 µl of 100% DMS (∼200 mM) was added to the refolded RNA and incubated for 1 minute in a thermoshaker (250 rpm and 37°C). The reaction was quenched with 40 µl of 100% 2-ME, pipette mixed, and placed on ice for purification.

### 2.5. In vivo dimethyl sulfate (DMS) probing of HeLa and THP1 cells

Growth phase cells were allowed to reach ∼80-90% confluence in 10 cm dishes (HeLa) and ∼900,000 cells/mL in T75 flasks (THP1). Cells were probed using a modified DMS-MaPseq protocol [29]. A 2% (v/v) DMS solution was freshly prepared before treatment of each dish using 25% ethanol and 75% DPBS. For HeLa cells, growth media was removed from the cells before treating with 3 mL of DMS solution for 1 minute at room temperature with agitation. The DMS was removed before cells were neutralized twice with 4.5 mL of dithiothreitol (DTT; Sigma-Aldrich) in DPBS (5X molar excess to DMS). Cells were harvested in 1 mL of TRIzol® (Invitrogen). For THP1 cells, 1.0x10^7^ cells per reaction were pelleted at 200*xg* for 3 minutes (for all subsequent unless otherwise stated) after cell counting (hemacytometer). Growth media was aspirated, cells were washed with 5 mL of DPBS before pelleting. Cells were resuspended in 1 mL of 2% DMS solution mentioned above and incubated for 1 minute with agitation at room temperature. The reaction was quenched by pipette mixing in 1 mL of DTT in DPBS (5X molar excess to DMS), followed by cell pelleting. The neutralized DMS was removed, and the cells were subjected to 1 mL of TRIzol® (Invitrogen). All DMS probing of cells was completed in duplicate.

### 2.6. Isolation and purification of DMS probed RNA

Both in vitro and in vivo DMS probed RNA was isolated using a minimally modified Direct-zol Plus RNA mini-prep kit protocol (Zymo). Briefly, 200 μL of 1-bromo-3-chloropropane (Sigma-Aldrich) was added to the sample and shaken vigorously. After a 2-minute room temperature incubation, samples were centrifuged at 12,000*xg* for 15 minutes at 4°C. The upper aqueous phase of each sample was transferred to new 1.5 mL microcentrifuge tubes and an equal volume (∼500 μL) of 100% ethanol was added. After pipette mixing the ethanol, each sample was loaded onto Direct-zol Plus RNA mini-prep columns, and all downstream DNase treatments and washes were done according to manufacturer’s protocol (Zymo). After elution, a NanoDrop One (Thermo-Fisher) spectrum was taken to obtain the concentration of total RNA.

### 2.7. Induro reverse transcription, PCR amplification, and quality control

Purified RNA was used for reverse transcription (RT) with the Induro enzyme (NEB) following the manufacturer’s protocol with 20 μL reactions using 1 µg of RNA. For in vitro probed RNA, random hexamer primers (IDT) were used, and for in vivo probed RNA a 1:10 ratio of poly dT (Invitrogen) and random hexamer (IDT) primers were used. Reactions were incubated at 25°C for 2 minutes, 55°C for 30 minutes, and 95°C for 1 minute. RT products were diluted to 10 ng/μL for PCR amplification. Targeted amplification of the *TNFRSF1A* 3’UTR was performed on the cDNA. PCR was completed using 20 μL Q5 DNA polymerase (NEB) reactions. For in vitro probed samples, a single primer set was used to amplify the 386 nt, 836 nt, and 745 nt fragments of the TNFR 3’UTR s7, TNFR 3’UTR, and TNFR 3’UTR d7 constructs respectively (Fwd: 5’-CAGGCACCTGTTTCGGATTCG-3’ and Rev: 5’-GAACGAACGCATGTTGAACCGG-3’). For in cell probed samples, a single primer set was used to amplify a 739 nt fragment of the TNFR CDS and 3’UTR from HeLa cells (Fwd: 5’-GAGCGACCACGAGATCGATCG-3’ and Rev: 5’-CTTCAGCTGGAGCTGTGGACT-3’), and a single primer set was used to amplify 693 nt fragment of the TNFR CDS and 3’UTR from THP1 cells (Fwd: 5’-GAGCGACCACGAGATCGATCG-3’ and Rev: 5’-CATAGCAAGCTGAACTGTCCTAAGGC-3’). For HeLa and THP1 the last 31 nt and 77 nt of the 3’UTR could not be amplified, respectively. All PCR products sizes were validated via agarose gel (**S2 File**) before determining their concentration on a Qubit® 2.0 Fluorometer (Thermo-Fisher).

### 2.8. Library preparation, quality control, and sequencing

All PCR products were put through Illumina DNA library preparation following the manufacturer’s protocol. Input DNA concentrations were 150 ng and 324 ng for HeLa and THP1 in vivo probed samples, respectively, and 25 ng for all in vitro probed samples. Briefly, DNA was tagged with adaptor sequences and fragmented using Illumina’s tagmentation process. Tagmentation products were cleaned up and amplified with Illumina Nextera DNA UD Indexes (IDT) following Illumina’s recommended PCR parameters. Resulting libraries were cleaned following the standard DNA input protocol. Resulting library concentrations were determined using the Qubit® 2.0 Fluorometer and library sizes were determined using an Agilent 2100 Bioanalyzer. All in vitro libraries were determined to be between 300-600 nt and all in vivo libraries were determined to be between 500-600 nt (**S2 File**). All sequencing was performed on an iSeq100 benchtop sequencer (Illumina) using paired-end reads (150 x 150 nt). Following the manufacturer’s suggestions, all libraries were diluted to 1 nM with Illumina Resuspension Buffer (RSB) before combining all in vitro libraries together and all in vivo libraries together at equal volumes. Following the manufacturer’s suggestion, the pooled libraries were further diluted to 100 pM with RSB before loading 20 μL into the sequencing cartridge and starting the sequencing run.

### 2.9. Analysis of sequencing data with RNA Framework and DRACO

All sequencing reads were output as fastq files that were then analyzed using the `RNA Framework` package [30, 31] for analysis of next generation sequencing data generated by RNA structure probing experiments. The fastqc program was first used to determine phred (quality) scores and adaptor content. The cutadapt function was used to trim off any adaptor sequence before reanalyzing with fastqc. Trimmed fastq files were indexed with Bowtie2 [32, 33] for read mapping using the known in vitro construct sequences and the *TNFRSF1A* MANE (ENST00000162749.7) fasta file. All paired end reads were mapped to the corresponding fasta files using the “rf-map” function. All mapped bam files from the same library were merged to allow for both individual and combined processing. Coverage and read depth information was obtained using SAMtools on the bam files. The “rf-count” function was used to determine the frequency of nucleotide mutations and a rc file was generated. Using the “rf-norm” function, mutations for A and C nucleotides were normalized and an xml file of reactivities was generated. The “rf-fold” function was used to fold all xml files. Here, the –sh, -dp, and -md flags were used for reactivity informed 120 nt and 600 nt max base pair models (output as ct files) as well as Shannon entropy and base pair probability files. Use of `DRACO` [34] determined whether dynamic regions were present. The mm file generated during the rf-count step, was used to make a json file for further processing. The json file and previous rc files were used as input in the “rf-json2rc” function to generate a `DRACO` rc file, which was then processed by the “rf-norm” function to generate xml files for each unique reactivity profile. The “rf-fold” function was used to fold all `DRACO` xml files and produce individual ct files. Using the python script “xml_reactivity_full_extract_batch.py” all xml files were converted to react files (folding constraints) for `ScanFold`. Additionally, the react files were further converted to heat maps for visualization of reactivities on 2D structure models in `VARNA` [35]. For more information on `RNA Framework` visit (https://rnaframework-docs.readthedocs.io/en/latest/) and for `DRACO` visit (https://github.com/dincarnato/DRACO).

### 2.10. Covariation analysis

All structures with z-scores ≤ -2 were analyzed for covariation using the cm-builder Perl script [31]. This script utilizes `Infernal` (release 1.1.2) [36] to build and find covariance models for `ScanFold` predicted structures using large sequence databases. `Infernal` databases were created using an NCBI `BLAST` search of the *TNFRSF1A* transcript ENST00000162749.7. Here, the NCBI Refseq database was searched using the following parameters: `BLAST`n, gap open 5, gap extend 2, reward 1, penalty -1, max target sequences of 5000. All pseudogenes, “like” sequences, and non-*TNFRSF1A* sequences were unselected before the resulting list was downloaded and used. The structural alignment files from `Infernal` (in Stockholm format) were tested for covarying base pairs and analyzed with both `CaCoFold` and `R-scape` (version 1.5.16 with `ScanFold` predicted structures). Statistical significance was evaluated by the APC corrected G-test [37, 38] using the default E-value of 0.05. Power files were generated and analyzed using an in-house script that bins the power of covarying base pairs into 0 - 0.1, 0.1 - 0.25, and > 0.25. All input files, Stockholm alignments, `R-scape`/`CaCoFold` results, and power analysis data can be found in **S3 File**.

### 2.11. Reporter plasmid design and cloning

All plasmid constructs were based on the `ScanFold` predicted secondary structures; exact regions were determined based on structure and miRNA binding site proximity. All RNA tested was cloned from the 3’UTR of *TNFRSF1A* (MANE) into an intron modified pmirGLO plasmid (Promega) designated with an “i” (pmirGLOi) [20]. These sequences were ordered as gBlocks or Ultramers (IDT) for cloning. Cloning was completed using the HiFi Assembly kit (NEB) with HiFi reactions run at 50°C for 30 minutes, followed by transformation in NEB-5*α* competent *E. coli*. For HiFi cloning, a minimum of 15 nucleotides complementary to the upstream and downstream sequence surrounding the XhoI restriction enzyme site of the vector were added to each end of the sequences. When necessary, PCR was employed for amplification or the addition of homology arms. Colony PCR was used to determine insertion, and sequences were validated by Sanger sequencing (Iowa State DNA Facility). A schematic of the plasmid, and information regarding construction, the cloning strategy, and primers used can be found in **S4 File**.

### 2.12. Dual luciferase assays and qPCR for translational efficiency

HeLa cells were trypsinized from a 10 cm dish at 80-90% confluence. Cells were counted using a hemacytometer and plated at 2x10^4^ cells/well in a 96-well dish (6 biological replicates per construct per experiment) for dual luciferase assays and at 1x10^5^ cells/well in a 24-well dish (3 biological replicates per construct per experiment) for qPCR. Cells were transfected 24 hours later with experimental or control plasmids (empty pmirGLOi or containing the whole *TNFRSF1A* 3’UTR) using Lipofectamine 3000 (Invitrogen). Respectively, 5 or 25 ng of dual luciferase plasmid was transfected into each well of the 96- and 24-well plates with 95 or 475 ng of pUC19 plasmid filler, respectively. Cells were supplemented with fresh DMEM 24 hours post-transfection and analyzed (dual luciferase assay) or harvested (for RNA) 48 hours post-transfection.

A Promega Dual Luciferase kit was used following the manufacturer’s protocol. In brief, cells in the 96-well dish were washed with DPBS before lysing with 1x passive lysis buffer (PLB). Lysate from each well was transferred to an opaque white 96-well dish for recording luminescence using a dual injecting GloMax Explorer (Promega). The Relative Response Ratio (RRR) was calculated by dividing the light units of Firefly luciferase by those of Renilla luciferase on a per-well basis. The RRR was normalized to the appropriate controls per experiment to account for experiment-to-experiment variability. Resulting values were plotted as the mean ± standard deviation. Using a two-tailed, equal variance T-test, the significance of changes was set at a p–value of < 0.05 from a minimum of three biological replicates per condition (**S5 File**).

RNA was isolated from the 24-well dish cells using 400 μL of TRIzol® (Invitrogen) per well and the same modified Direct-Zol RNA mini-prep (Zymo) protocol used in the isolation of DMS probed RNA above. cDNA was generated following the Superscript III (Invitrogen) RT manufacturer’s protocol. Here, between 200 ng and 1 μg of RNA was used with random hexamer (IDT) priming. For quality control, no RT controls were completed for all samples. All cDNA samples were diluted with water to ∼5 ng/μL based on RNA input and qPCR was performed with 1 μL of diluted cDNA. Reactions were multiplexed with cPrimeTime® primer/probes (IDT) designed to overlap the introduced intron for each of the Firefly and Renilla luciferase genes (Firefly: forward 5′–ACAAAACCATCGCCCTGATC–3′, reverse 5′–ATCTGGTTGCCGAAGATGG–3′, probe 5′6-FAM / ACCGCTTGT / ZEN / GTCCGATTCAGTCAT / 3′IABkFQ; Renilla: forward 5′–CCTACGAGCACCAAGACAAG– 3′, reverse 5′–ACCATTTTCTCGCCCTCTTC–3′, probe 5′SUN / CACGTCCAC / ZEN / GACACTCTCAGCAT / 3′IABkFQ) and run using PrimeTime® Gene Expression Master Mix (IDT) on a QuantStudio3 (Thermo-Fisher). Ct values were calculated using the automatic threshold detection settings of the QuantStudio Design & Analysis desktop software (v1.5.1). The dCt (to Renilla) was calculated on a per well basis. The dCt was used to determine fold change (2^-ΔCT^). Translational efficiencies were calculated by dividing the RRR by the mRNA fold change. Maximal data variance was maintained per condition by using the mean of whichever set (RRR or mRNA) had lower variance per experiment. Normalization was done per experiment to the appropriate control (empty vector or whole 3’UTR, as indicated) before combining all data. Outliers for translational efficiencies were removed using Grubbs’ Test for outliers. Using Student’s T-test with a p-value of < 0.05 for significance (two-tailed, equal variance), experimental constructs were compared to the appropriate control (**S5 File**).

### 2.13. Biotinylated-RNA pulldowns, mass spectrometry, and immunoblotting

Preparation of HeLa and THP1 whole-cell lysates began with cell suspension (HeLa trypsinized), counting, and centrifugation for 3 min at 200*xg*. Cell pellets were washed once with ice cold DPBS, pelleted again, and resuspended at 10^7^ c/mL in ice cold polysome extraction buffer (PEB: 20 mM Tris, pH 7.5; 100 mM KCl; 5 mM MgCl_2_; 0.5% NP-40) [39] containing 1 mM PMSF, HALT, 200 U/mL RNaseOUT (Thermo-Fisher). The resuspended pellet was incubated on ice for 15 min prior to Dounce homogenization. The homogenates were spun at 15,000*xg* for 10 min at 4°C for collection of the supernatant. Lysates were pre-cleared using streptavidin-agarose beads (Thermo-Fisher) and concentrations were determined using a BCA assay (Thermo-Fisher).

To generate biotinylated RNA, plasmids containing *TNFRSF1A* regions were used. DNA templates containing T7 recognition sequences were made by PCR amplification (Q5, NEB) for the full-length 3’UTR, deletion of structure 7 (d7), just structure 7 (s7), and the reverse complement (“rc”) of structure 7 (negative control). T7 recognition sequences were incorporated into the primers found in **S6 File**. PCR products were *in vitro* transcribed with T7 RNA polymerase (Invitrogen) and biotinylated CTP (1 in 4) to generate biotinylated RNA. RNA was NanoDrop® quantified and purified using both TRIzol® and a Zymo Direct-zol column prior to size validation by National Diagnostics urea-PAGE as previously.

Precipitations were done using 1 ug of biotinylated-RNA and 0.5 mg lysate in a Tris-EDTA-NaCl-TritonX-100 buffer (TENT: 10 mM Tris pH 8.0, 1 mM EDTA pH 8.0, 250 mM NaCl, 0.5% Triton X-100). Mixture was incubated for 30 minutes at room temperature both before and after addition of buffer-equilibrated streptavidin agarose beads (∼50% slurry). Beads were collected by centrifugation at 7000*xg* for 30 seconds at 4°C and washed four times with 1 mL ice-cold TENT buffer before elution with 2x Laemmli buffer. Samples were run on a 4-15% SDS PAGE gel (Bio-Rad) before silver staining (Bio-Rad). Unique bands were excised from the gel and LC-MS/MS analysis was performed (Iowa State Protein Facility; **S6 File**). Mass spectrometry results were searched against only human protein data in the UniProt database. This was performed once for each cell type lysate on different days but were combined for purposes of immunoblotting. To verify HuR (*ELAVL1*) binding to structure 7, some sample was subjected to SDS-PAGE and transferred to PVDF membrane before immunoblotting (Santa Cruz Biotechnology, sc-5261).

### 2.14. RNA immunoprecipitation

All RNA immunoprecipitations were similarly done as before [40]. Briefly, dishes of HeLa, Lenti-X, and THP1 cells were rinsed twice with 10 mL of cold DPBS before lysis in 500 μL (per two 10 cm dishes) of ice-cold RIPA lysis buffer (50 mM TrisHCl; pH 8.0; 1 mM EDTA; 150 mM NaCl; 100 μM Na_3_VO_4_; 1.0% NP-40; 1.0% sodium deoxycholate; 0.1% SDS) containing protease/RNase inhibitors (1:100 HALT, ThermoFisher; 1 mM PMSF, Sigma-Aldrich; 300 U/mL RNaseOUT, Invitrogen). Lysates were sonicated on ice four times at 50% amplitude for 5 seconds each prior to centrifugation to clear the lysate (15,000*xg*, 10 min, 4°C). The supernatant was diluted 2.5-fold with detergent-free lysis buffer containing 200 U/mL of RNaseOUT, followed by pre-clearing with 100 μL of protein A/G-PLUS agarose beads (Santa Cruz Biotechnology) for 1 hr at 4°C. Lysates were incubated with either anti-HuR (sc-5261) or mouse IgG antibody (sc-2025) for 1 hr at 4°C on the rotisserie. Beads were then added for an additional hour at 4°C. For input samples, 25% of the volume added for each antibody of the pre-cleared lysate was set aside for RNA isolation. Beads were washed three times with cold DPBS (1 mL each) before TRIzol® was added (to the input as well). Samples were incubated for 15 minutes at room temperature prior to either storage at -80°C or completion of RNA isolation. Total RNA isolation was done using a Zymo RNA Clean & Concentrate kit per manufacturer’s protocol. Input lysate was eluted at a higher volume to make the input equivalent to 10% of the immunoprecipitation samples for reverse transcription (RT). No RT controls were done using the input lysate. Total RNA, random hexamers (IDT), and Superscript III (Thermo-Fisher) were used for RT. qPCR over the structure 7 region (forward 5′–GTTCGTCCCTGAGCCTTT–3′, reverse 5′– ATAGAAACTTGGCACTCCTGT– 3′) was used to determine enrichment relative to input. This was done one time in each of the three different cell types.

## 3. Results

### 3.1. Lowest ScanFold z-score regions favor non-coding RNA in the *TNFRSF1A* gene and transcript

To predict RNA regions with structural and functional propensity, `ScanFold` was used to analyze the 13,357 nucleotide *TNFSFR1A* pre-mRNA (ENSG00000067182), as shown in **Fig 1A**. `ScanFold` predicts many sequence regions with z-scores ≤ -2 throughout the entire pre-mRNA. Almost all the predicted lowest z-score base pairs occurred in untranslated regions (UTRs) or introns. The exception to this could be found in the penultimate exon, which encodes the neutral sphingomyelinase activation domain (NSD). ENSEMBL predicts that the pre-mRNA can be spliced into 18 different isoforms, and of these isoforms, 12 encode proteins (seven of which are subject to nonsense mediated decay) and six are processed but do not produce protein. These transcripts range in size from 571 to 2,379 nucleotides. NCBI predicts four different forms, three of which are protein coding. We focused on the MANE (Matched Annotation between NCBI and ENSEMBL) transcript isoform (ENST00000162749 or NM_001065.4), which represents a high-quality reference transcript. `ScanFold` predicted only four moderately stable (comprised of base pairs with average z-scores between -2 and -1) structures throughout the 5’UTR and coding sequence of the mRNA and four very small hairpin structures with z-scores ≤ -2 (**Fig 1B**). There were, however, six low z-score regions in the 3’UTR, suggesting that the 3’UTR of *TNFSFR1A* has the greatest propensity for ordered stability and structure over other regions of the transcript. This finding is also congruent with the known regulatory roles of 3’UTRs [41], so we conducted a more focused analysis of the 3’UTR to evaluate the activity of the predicted RNA structures.

**Figure 1.**
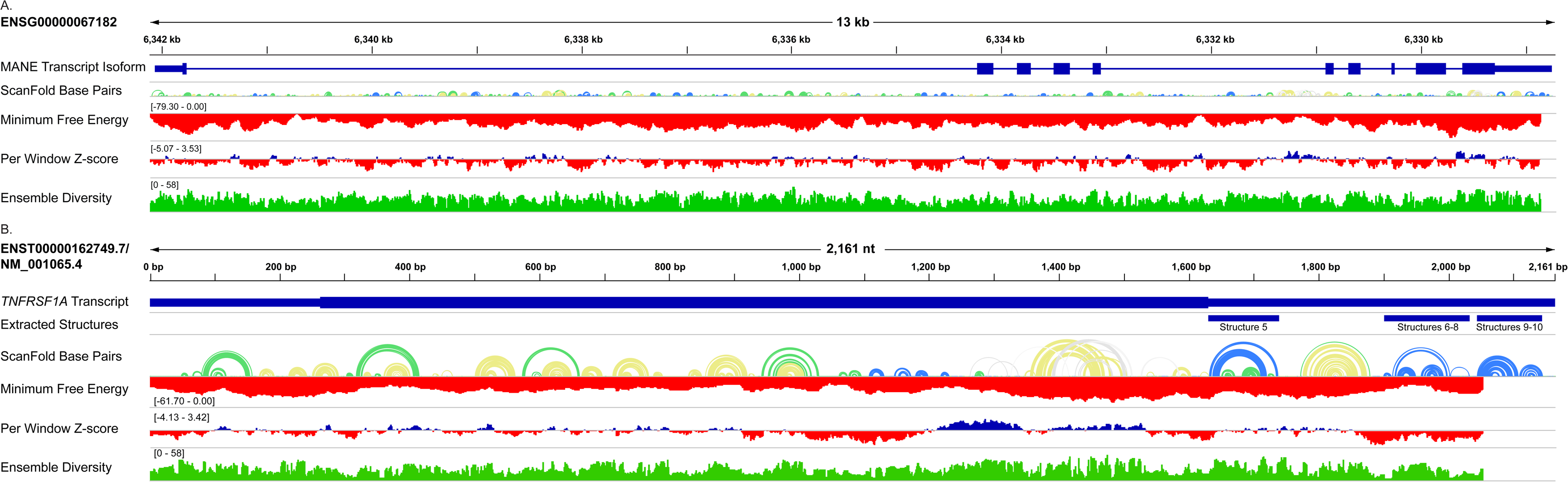
`ScanFold` output for *TNFRSF1A* gene (**A**) and transcript (**B**). Data presented as IGV tracks includes forward window-based calculations of the minimum free energy, z-score, and ensemble diversity (numbers represent low and high values), as well as the z-score-weighted base pairing (arc color represents z-score > 0, >–1, >–2, and ≤-2 color in white/gray, yellow, green, and blue, respectively). The gene schematic in **A** shows the introns (line) and exons (taller boxes), with demarcation between coding (tallest) and untranslated regions, from the MANE transcript. **B** shows a representation of this transcript in its spliced form and includes a track for extracted structured regions.

### 3.2. The *TNFRSF1A* 3’UTR affects translational efficiency

To determine whether any of the low z-score regions identified in the 3’UTR were functional, we cloned the full-length 3’UTR (541 nt), fragments that encompassed halves of the 3’UTR (3u5e, 250 nt; 3u3e, 291 nt), and predicted low z-score structured regions either in isolation or that were close in proximity or included predicted miRNA binding sites (structures [s]5, 108 nt; s6-8, 132 nt; s7, 90 nt; s9-10, 100 nt; s9end, 139 nt; **Fig 2A**), into custom dual luciferase reporter assay plasmids. Translational efficiency was then assessed in HeLa cells following protocols we previously established for other gene targets (see **S5 file** for schematic) [20, 21]. All of the constructs exhibited significantly reduced mRNA as well as relative response ratio or protein to the empty vector control (Vec). The full-length 3’UTR of *TNFRSF1A* inhibited translational efficiency as shown in **Fig 2B** (bottom). To a lesser extent, but still significantly, each of the 5’end or the 3’end alone (3u3e or 3u5e, respectively) also impaired translational efficiency. This latter effect was also mimicked by a 139 nt section at the 3’ end of the 3’UTR (s9end) that retained a `TargetScan` predicted binding site for miR-142-3p. When the predicted miRNA-binding site was not present (s9-10), the translational efficiency was statistically indistinguishable from the empty vector. The presence of another miRNA binding site, both predicted and verified previously (miR-29a-3p; [23]), just downstream of structure 5 (in 3u5e) also coincided with reduced translational efficiency, as did s5 alone, which only had two samples in the analysis. Comparing s6-8 with s7 indicated that a second predicted binding site for miR-142-3p did not affect translational efficiency in our assays. Taken together, these data implicate the sequence region just downstream of structure 10 and as well as the region in and around structure 5 as important for reduced translational efficiency of the *TNFRSF1A* 3’UTR. Notably, the regions with predicted miRNA binding sites were also predicted to be unstructured or looped out and accessible by `ScanFold` (**Fig 2A**).

**Figure 2.**
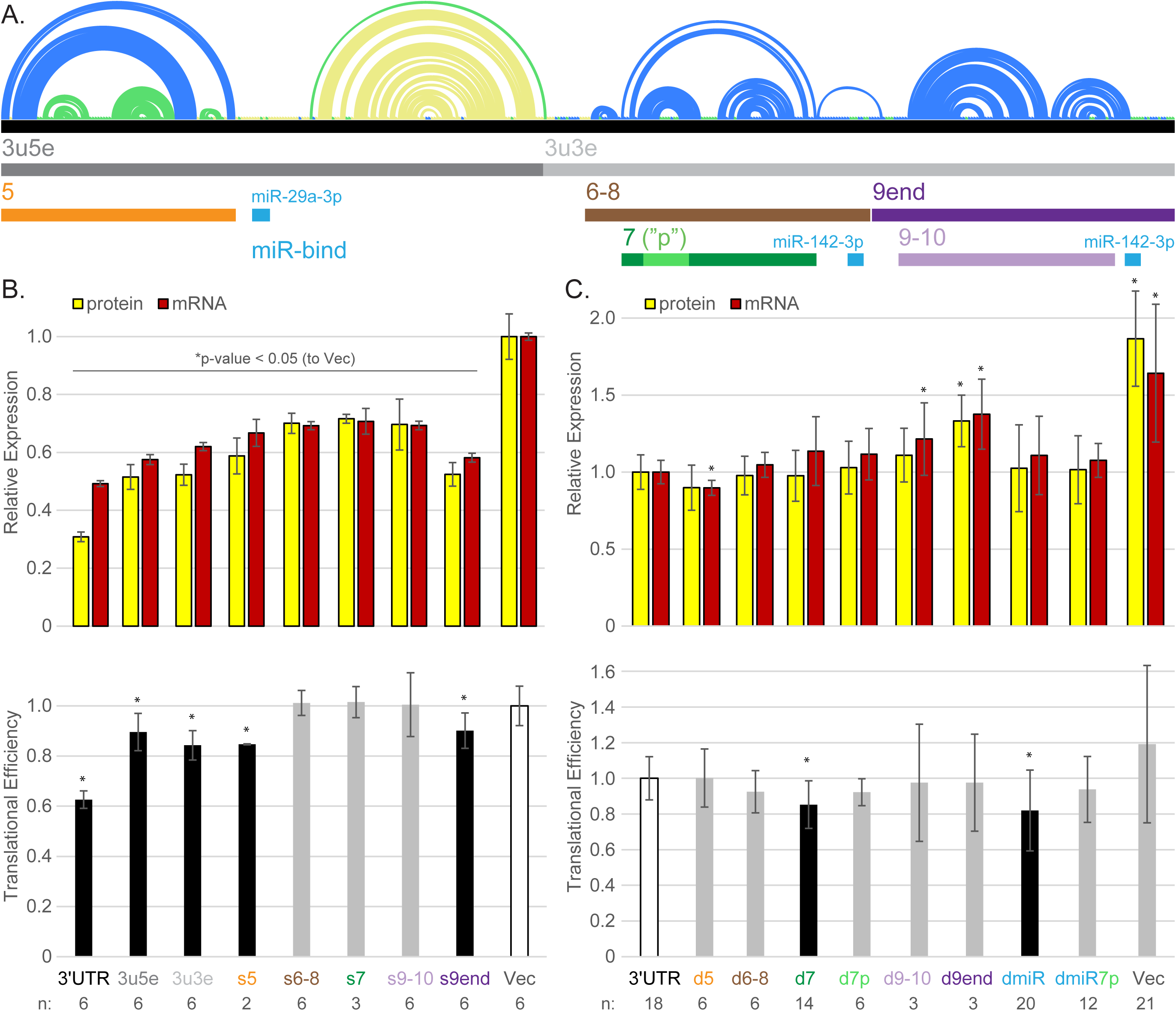
Identification of regions in the 3’UTR of *TNFRSF1A* that alter translational efficiency. **A.** Schematic of the 3’UTR of *TNFRSF1A* showing the `ScanFold`-predicted base pairing (color scheme as in Fig 1). Colored bars below represent regions that were cloned into a dual luciferase reporter construct (see S5 File for backbone) or deleted from the 3’UTR. For the grey bars, the “3u” designation indicates the 3’UTR whereas the “5e” or “3e” indicates the 5’ or 3’ ends, respectively. The numbers represent the `ScanFold`-identified ≤ -2 structured region that is encompassed. miR-bind regions are those predicted by `TargetScan`. **B.** The top portion shows reporter assay results for the relative level of protein (by luminescence; gold) and mRNA (by qPCR; red) after normalization to both a transfection control (Renilla) and empty vector (Vec). Translational efficiency, or protein per mRNA, is shown below. The number (n) of independent data points included per condition is given. The indicated conditions-what was expressed from the 3’UTR-are provided at the bottom. **C.** Reporter assay results, like B but with notable differences. These are deletion (“d”) constructs where only the indicated region has been removed from the 3’UTR, and these were normalized to the full-length 3’UTR. For both B and C, both asterisks and black bars represent statistically significant (p-value < 0.05; t-test, equal variance, two-tailed) changes from Vec (B) and full-length 3’UTR (C), respectively.

Along with testing sufficiency, the necessity of `ScanFold`-identified 3’UTR regions to alter translation was examined using deletion (“d”) constructs (**Fig 2C**). Compared to the full-length *TNFRSF1A* 3’UTR, d5, d9-10, d9end, and Vec had significantly different levels of mRNA. These data suggest that region 5 may have a stabilizing whereas d9-10 and d9end may have a destabilizing effect on the transcript. Only d9end and Vec also had correspondingly significant changes in relative protein. Experiments with Vec did not reach statistical significance compared to the 3’UTR in this set of experiments. Removing all three of the predicted miRNA binding sites (dmiR) not only did not enhance translation efficiency compared to the entire 3’UTR, it subtly reduced it. Removal of structure 7 (d7) also showed a modest but significant reduction in translational efficiency, suggesting that structure 7 is important for optimal translational efficiency.

### 3.3. Analysis of evolutionary conservation and covariation highlights the importance of ScanFold-identified 3’UTR regions

Structures identified in the 3’UTR already garnered our attention from the `ScanFold` analysis because of their highly negative z-scores. The analysis of the evolutionary conservation of RNA structure predicted in *TNFRSF1A* further reinforced this finding. Structures 5, 7, and 9 are supported by at least one covarying base pair (**S3 File**). A co-mutation to preserve structured pairing is not typically a random event, but rather indicates the importance of maintaining a structure for function and serves as a first-pass validation of model base pairing [42, 43]. One covarying base pair was identified in structure 7, shown in **Fig 3A** (grey shading) in the first hairpin of the multi-branched `ScanFold` model. This covarying base pair appeared in both the `ScanFold` structure-modeled analysis (`Rscape`) and the evolutionarily modeled analysis (`CaCoFold`; **S3 File**). In addition to conservation of secondary structure, the sequence of structure 7 was highly conserved (72.7% APSI) across mammals (**Fig 3B**). Notable were two distinct conserved poly-uridine (poly(U)) tracts: one well-defined (U)_5_ tract between the two `ScanFold` identified hairpins and a second, longer, U-rich region with more variable length and composition, which encompasses the 3’ half of the second hairpin (pairing with an upstream A-rich stretch of sequence). The presence of these conserved U-rich segments suggested possible docking sites for various poly(U) binding proteins, which are known to play important roles in post-transcriptional gene regulation [44].

**Figure 3.**
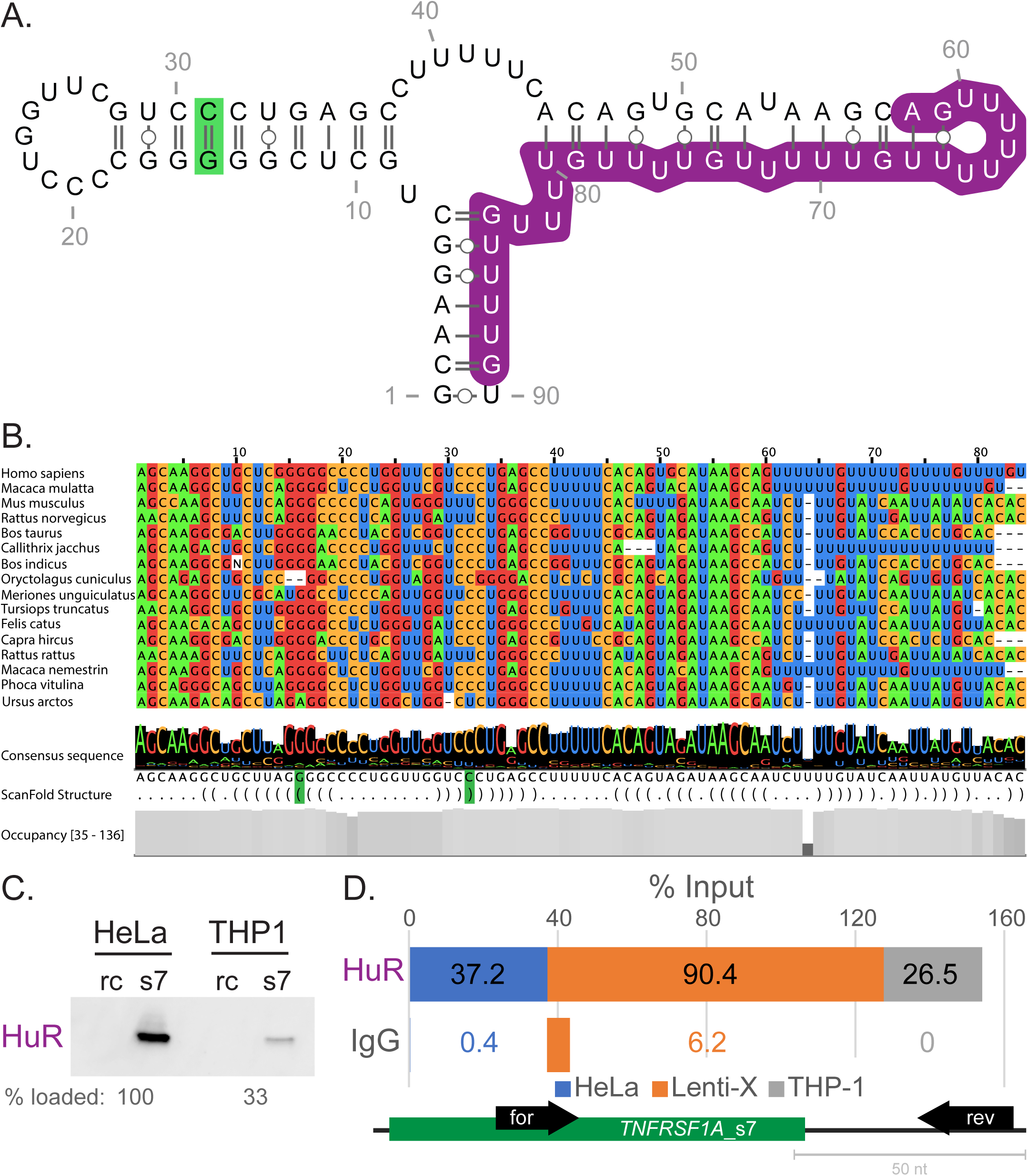
*TNFRSF1A* structure 7 (s7) is conserved and is associated with HuR. **A.** `ScanFold` s7 model showing the RBPmap-predicted binding region for HuR binding (purple). The covarying pair is indicated with green shading. **B.** *TNFRSF1A* s7 is conserved across mammals. Representative sequence sub-alignment of 15 homologous sequences for s7 (from a total of 134 homologous sequences identified). Aligned nucleotides are colored according to identity and are followed by a track showing the consensus sequence data across all 135 mammalian species considered. This is followed by an aligned “dot-bracket” structure that corresponds to the 2D structure represented in A, with the covarying G-C base pair in the first hairpin highlighted green (note the variation in the alignment that yields “consistent mutations” G-C to G-U and “compensatory mutation” from G-C to A-U). The final track indicates nucleotide occupancy at each aligned site, which ranges from 35 to 136 (no gaps). **C.** Immunoblot results for HuR following biotin pulldown assays using the indicated cell lysates and either s7 or the reverse complement (rc) of s7. Less THP1 sample remained after mass spectrometry, accounting for the unequal loading between cell types. **D.** Stacked bar graph showing collective RNA immunoprecipitation results as a percentage of input. An antibody that binds HuR or an immunoglobulin isotype control (IgG) from the indicated cell lines were used for immunoprecipitation prior to RNA isolation and RT-qPCR for the region encompassing s7. Schematic at the bottom indicates the relative position of the forward (for) and reverse (rev) primers that were used for detection of s7 (darker green). In graph numbers indicate calculated percent input values. Experiments in B and C were performed once in each of the indicated cell types.

### 3.4. Identification of *TNFRSF1A* structure 7 RNA binding proteins

To begin to discover what proteins might be binding to structure 7 (s7), we performed biotin-pulldown assays with s7 as bait followed by silver-staining and mass-spectrometry of excised bands. This was done using both HeLa and THP1 cell lysates. Only two proteins were identified that were consistent between the two cell types, HuR (encoded by the gene *ELAVL1*) and RPS3A (**S6 File**). The findings coincided with known consensus ribonucleotide binding sequences of each protein: “NNUUNNUUU” for HuR and “YYYYTTYC” for RPS3A [45, 46]. This suggests that HuR would bind the second hairpin of structure 7, while RPS3A would bind the single-stranded (U)_5_ motif between the two branches (**Fig 3A**). Given the known role of HuR to promote translation [47], we supported the association for HuR through immunoblotting after biotin-pulldown (**Fig 3C**). RNA immunoprecipitation (RIP) from three different cell types also reinforced this association (**Fig 3D**). Notably, while these assays are not quantifiable because they were only performed once per lysate, they were done in different cell types, providing two and three biological replicates for the immunoblot and RIP, respectively. The consistent observation of HuR association across different cellular contexts (HeLa, Lenti-X, and THP1) suggests a reproducible interaction. Future quantitative and direct binding assays (e.g., EMSA) would be necessary to confirm the directness and strength of this interaction.

### 3.5. *TNFRSF1A* 3’UTR DMS-MaPseq supports ScanFold-identified structured regions

To validate the `ScanFold`-predicted secondary structure of the *TNFRSF1A* 3’UTR, particularly structure 7, we performed targeted DMS-MaPseq experiments—following protocols optimized in our previous work [21]. In-cell RNA structure probing was used to investigate the entire 3’UTR in both HeLa and THP1 cells, providing the first targeted structure probing for this transcript (**Fig 4**). We also performed in vitro probing with several constructs, including the entire 3’UTR, the 3’UTR with structure 7 removed, or segment 7 alone (**S2 Fig**). Both in vitro and in cell probing strongly supported the `ScanFold`-predicted structure 7, with two hairpins consistently identified across all constructs (**Fig 4B**; **S3 Fig**). The second poly(U) containing hairpin, however, could adopt an alternative hairpin conformation in the model developed using in vitro probing data from the 3’UTR. For structure 5, while in vitro probing agreed with `ScanFold`’s multi-branch loop prediction, in cell probing showed greater variation in HeLa and THP1 cells, suggesting potentially dynamic behavior. In contrast to the differential modeling observed in segments of structures 5 and 7, structure 9 was consistently formed across all in vitro and in cell models. These results highlight the ability of `ScanFold` to identify stable RNA structures and suggest certain regions within the *TNFRSF1A* 3’UTR that may exhibit dynamic folding in response to cellular conditions.

**Figure 4.**
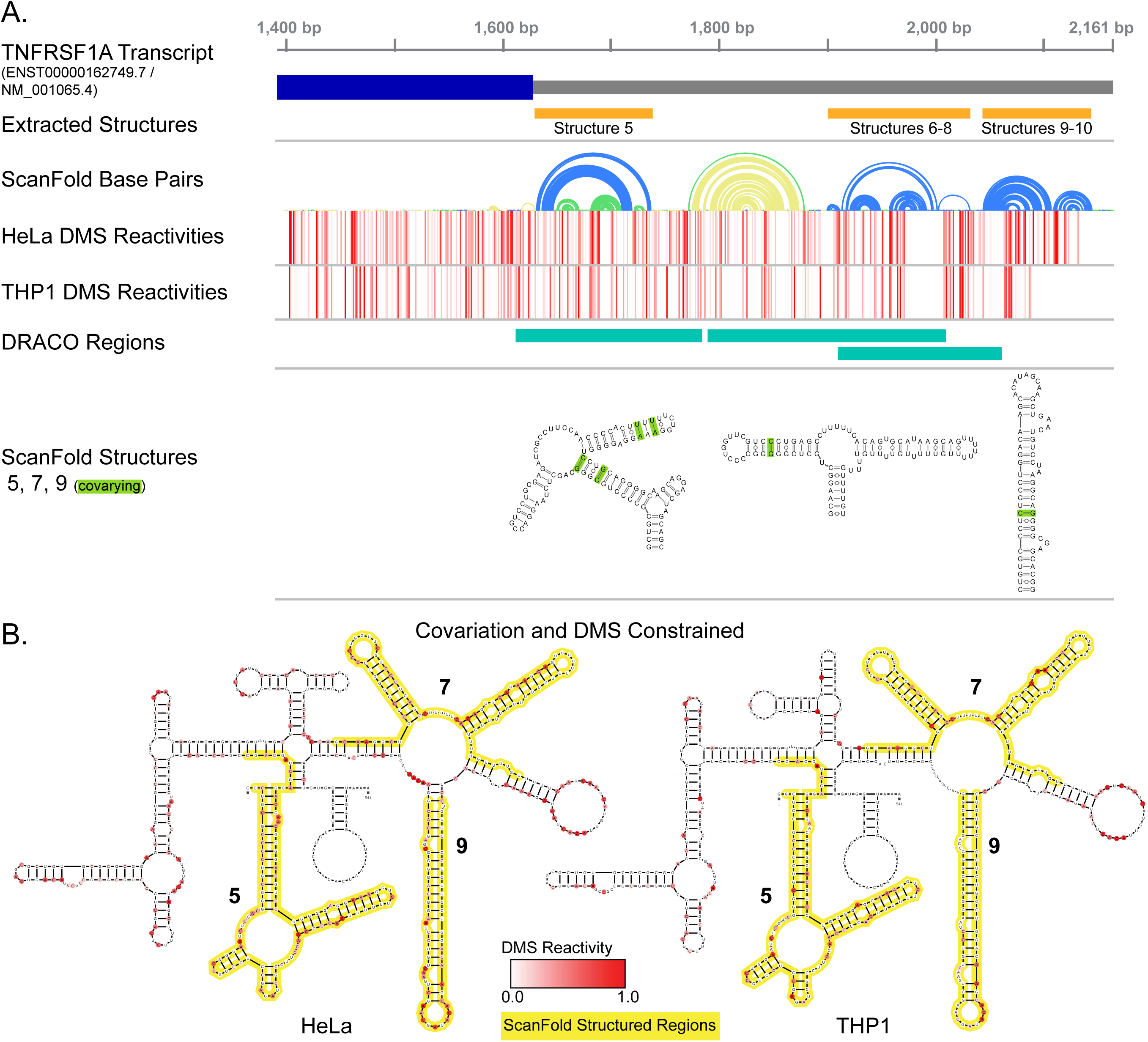
Targeted DMS-MaPseq structure probing supports `ScanFold` models. **A.** IGV tracks showing the region of the TNFRSF1A transcript covered by targeted structure probing (3’UTR is grey and a truncated coding sequence is dark blue). `ScanFold` predictions and extracted structures are as previously shown (Fig 1 and 2). DMS reactivities are shown in heatmap form for both HeLa and THP1 cells. `DRACO`-identified dynamic regions are marked in teal. Models for structures 5, 7, and 9, show the covarying base pairs (green). **B.** Informed `ScanFold` structure models from both HeLa and THP1 with hard constraints for covarying base pairs and soft constraints for DMS reactivities (please see Methods).

Given the potential for dynamic folding in the *TNFRSF1A* 3’UTR, we further explored possible conformational dynamics using `DRACO`. [34]. `DRACO` finds unique co-mutational events to extract unique reactivity profiles from MaPseq experiments, allowing for deconvolution of potentially dynamic structures. `DRACO` analysis of the in cell probing data revealed several potentially dynamic regions (**Fig 4A**). In THP1 cells, these regions fully encompassed structures 5 and 7 (**S4 Fig**). In HeLa cells, the dynamic regions within structure 7 started in the terminal loop of the first hairpin and extended past the second, poly(U) containing, hairpin (**S7 File**). Notably, the most dynamic region in structure 7 coincided with the HuR binding site and the poly(U) tract, suggesting a potential role for these elements in conformational changes. Overall, our analysis indicates that structure 5 and 7, or at least portions of their sequences, exhibit dynamic folding. This dynamic behavior may be important for the regulation of *TNFRSF1A* expression, potentially through interactions with RNA-binding proteins like HuR.

## 4. Discussion

Here, we computationally identified and experimentally characterized uniquely ordered RNA structures within TNFRSF1A that may contribute to post-transcriptional regulation. Significantly, the *TNFRSF1A* mRNA, encoding TNFR1, could be an important therapeutic target to mitigate inflammation, as intracellular signaling is critical to TNFR1-mediated outcomes. The complex signaling associated with this gene product and the fact that either over activation or under activation can lead to detrimental inflammation renders it challenging to target [48]. Pathological inflammation observed in TRAPS, which coincides with mutated TNFR1 that is thought not to be expressed on the cell surface [48, 49], further highlighting the importance of intracellular signaling. As a result, TRAPS-associated TNFR1 exhibit less TNFα responsiveness but display a curious increased intracellular signaling. This suggests that regulating the dose of the gene product could be a way to influence TNFR1-mediated outcomes.

Here we provided a first, small step in the process to understand the RNA structural landscape of *TNFRSF1A*. Focused on the 3’UTR, we found three structures (5, 7, 9) using both in vitro and in cell DMS-MaPseq (**Fig 4**). After testing their effects on gene expression, our modest effects *suggest* that structure 7 plays a role in optimal translation under the conditions we used (**Fig 2**). Whether this relates to the native transcript or to pathological conditions remains an open question.

We found that the RNA-binding protein HuR (encoded by *ELAVL1*) was associated with structure 7 in RNA pulldowns and RIPs (**Fig 3**). HuR serves as a context-specific RNA-regulator–known to stabilize RNAs [50]–and has well-documented roles in inflammation [51, 52]. HuR may be a candidate TNFR1 translational regulator through structure 7, indicated but not fully demonstrated here. Of note, HuR has been shown to bind near miRNA seed binding sites [15] to either antagonize or cooperate with miRNAs [53, 54]. A perfect seed match for miR-142-3p binding is located within 20-50 nucleotides downstream of the structure 7 HuR-binding region(s) (**Fig 2A**). Whether HuR antagonizes the binding of this or another miRNA in the context of *TNFRSF1A* mRNA remains to be tested. HuR has already been shown to contribute positively to TNF signaling through stabilization of TNFα mRNA [55]; indeed, others are working to identify or fabricate compounds that will inhibit its activity [56, 57]. The functional significance of HuR’s association with the mRNA of TNFR1 and whether it promotes TNFR1 translation, remain undetermined and requires further study. In the future, it will be interesting to see whether the expression level of TNFR1 is affected by inhibition of HuR and how that relates to TNFα-involved outcomes.

DMS MaPseq data provides additional circumstantial context for HuR’s association with structure 7 and speculation that it is through the second stem-loop branch of structure 7 [45]. Specifically, our probing results, performed under differing conditions and variable constructs (**Fig 5**), revealed potential alternative folds—when data were included as constraints and restraints in secondary structure prediction. Coupled with the `DRACO` results that indicate conformational dynamics in the second branch, our results provide additional insight into the stability and potential existence of RBP sites [58].

Specifically, structure 7 has the potential to fold into two distinct structures, where the poly(U) sequence containing second hairpin) may act as a “slippery” sequence—in this case, one that can shift base pairing between helices of similar thermodynamics (**S4 Fig**). Thus, and speculatively, accessibility could be gained through unfolding, by HuR binding that leads to a conformational shift, or by some temporal combination of the two.

Notably, irrespective of possible conformational dynamics in the second hairpin of structure 7, the first hairpin, which contains the covarying pair remains unchanged in structure across all conditions and constraints/restraints used in structure modeling. Deletion of the terminal end of the first stem-loop, however, was insufficient to reduce translational efficiency in our assays, further suggesting a role for the nucleotides of the second hairpin in the regulation of translation. Thermodynamic modeling of the d7p deletion mutant (**S5 Fig**) suggests that the possible HuR-binding hairpin would still form as well as an alternative upstream hairpin. What role, if any, the first stem loop or any structured element upstream of the possible HuR-binding region plays is still an open question. When analyzed by RBPmap [59], there were no RBPs identified with potential to bind this first hairpin sequence, though our unconfirmed mass spec results and data analysis suggest that the ribosomal protein RPS3A may bind the loop between the two branches [46]. It could be that this highly structured and deeply conserved hairpin serves to maintain the accessibility of this site; validation of this and whether this has any relevance to TNFR1 translation are all reserved to future studies.

The significance of structures 5 and 9 remain unknown, but it is worth noting that just because our reporter assays show modest or no significant changes in translational efficiency, does not mean that these structures are inert—other cellular contexts or conditions could yet reveal functions. With only a few samples, structure 5 seemed to modestly reduce translational efficiency. As for structure 9, a previously identified methylation of an adenosine in structure 9 increased the level of TNFR1 protein in esophageal squamous cell carcinoma cells without changing the level of the mRNA [20]. Clearly more studies in a range of contexts are needed to understand the role these structures might play in TNFR1 regulation. Global modeling of in vitro and in cell DMS data suggests that structure 5 folds into a multi-branch structure, almost identical to the in silico only `ScanFold` prediction, with the branches showing some evidence of dynamism. The similarity of conformations and the maintenance of the multi-branch nature of structure 5 may simply be the result of loops and bulges of the structure breathing, or possibly as the result of protein interactions, formation of a local pseudoknot, or long-range interactions. The most consistent DMS data came from structure 9, which folded into the `ScanFold` predicted structure regardless of constraints, had no evidence of dynamics, and showed very few reactivities that conflicted with predictions (**S3 Fig**).

Beyond the discreet structures defined by `ScanFold`, scanning window data provide additional useful results. For example, results across the *TNFRSF1A* transcript can facilitate the development of oligonucleotide-based approaches for therapeutic targeting or basic research. To clarify, scans define the stability landscape, so high free energy (less stable) regions that overlap regions of positive z-score (unusually unstructured) are likely more accessible for intermolecular interactions. `ScanFold`, in this way, may be used–though not directly tested in these studies–to identify sequence regions that favor accessibility. Indeed, miR-29a-3p, which was previously shown to decrease both *TNFRSF1A* mRNA and TNFR1 protein [60, 61], falls within such a predicted region. Consistent with previous studies, we observed both reduced mRNA and luciferase activity with a reporter that contained a seed binding site for miR-29a-3p (3u5e) in HeLa, a cell type that is known to express miR-29a [62, 63]. Ambiguity remains surrounding this, however, as structure 5 alone also resulted in the same. Our data suggest that the more distal of the predicted miR-142-3p sites (9end) might be active, though only very low levels to none of this miRNA is expressed in HeLa cells [63]. Counterintuitively, translational efficiency was reduced when all three miRNA seed binding regions were removed. While the reason for this is uncertain, co-removal of the first hairpin of structure 7 seems to dampen this effect (dmiR vs dmiR7p, Fig 2C). Regardless, our structural analyses indicate accessible sequences for these miRNA sites (**S6 Fig**). Interestingly, viral 5’-isomiRs of miR-142-3p have been identified in several viruses, including human herpes virus 8 (HHV8), which can cause human Kaposi’s sarcoma [64]. A wild speculation could be that a viral-mediated reduction in the level of TNFR1 could inhibit important inflammatory recruitment, possibly permitting abnormal/unchecked endothelial proliferation in certain contexts. Circumstantial support for this is that Kaposi’s sarcoma is a rare side effect after treatment with anti-TNF-α therapy [65].

While this study provides novel insights into the RNA structural landscape of the *TNFRSF1A* 3’UTR, certain limitations warrant consideration. Although our findings highlight the role of structure 7 and its potential interaction with HuR, expanding studies to diverse cell types, such as disease-relevant models like TRAPS patient-derived cells, could reveal context-specific regulatory mechanisms and broaden the applicability of our results. The identification of miRNA binding sites (e.g., miR-29a-3p, miR-142-3p) within the context of 3’UTR structure opens promising avenues for future experiments to explore how RNA structure influences miRNA-mediated regulation of TNFR1 expression. Furthermore, investigating the dynamic folding of structures 5 and 7, particularly through mutational analysis of the poly(U) tract, will enhance our understanding of their regulatory roles. Excitingly, targeting structure 7 with innovative approaches, such as small molecules or antisense oligonucleotides, holds potential for developing novel therapeutic strategies to modulate TNFR1 expression in inflammatory diseases.

## 5. Conclusion

Our results identify multiple regions across *TNFRSF1A* with the ability to adopt unusually thermodynamically stable and ordered RNA structures. In addition to the 3’UTR structures that we highlighted in this study, the pre-mRNA sequence of *TNFRSF1A* is predicted to contain highly ordered and stable structures that overlap splice sites or fall within intronic sequences. Such structures could be playing roles in the modulation of splicing and may be amenable to structure/function analyses [66] or even drug targeting [58]. This work lays the foundation for future studies to definitively determine the functional impact of RNA structures on TNFR1 production and inflammation, and to explore their potential as therapeutic targets.

## SUPPORTING INFORMATION

Supporting information is available on Zenodo: https://zenodo.org/records/15692666

## DATA AVAILABILITY STATEMENT

All data are available in the manuscript and the Supporting Information.

## ACKNOWLEDGEMENTS

Thank you to the Iowa State University Research IT group for their support over the course of this project and members of the Moss Lab for their input.

